# *MALAT1* Levels Are Elevated in Thyroid Tumors from Chilean Patients with Lymphatic Infiltration and Are Linked to Metabolic Reprogramming

**DOI:** 10.64898/2025.12.23.696116

**Authors:** Michelle Tobar-Lara, Andrea Matamoros, Marcelo Muñoz-González, Diego Leiva, Gunther Redenz, Gino Nardocci, Luis Meneses, Patricio Cabané, Alvaro A. Elorza, Rodrigo Aguilar

## Abstract

Metastasis-associated lung adenocarcinoma transcript 1 (*MALAT1*) is a long non-coding RNA (lncRNA) implicated in cancer progression. In thyroid cancer, *MALAT1* has been proposed as a potential biomarker, but its role in disease progression remains incompletely understood. Here, we analyzed *MALAT1* RNA levels in paired tumoral and adjacent non-tumoral thyroid samples from a Chilean patient cohort. We found a positive correlation of *MALAT1* levels with lymphatic infiltration that was not replicated when modeling MALAT1 expression in a larger cohort obtained from the TCGA-THCA database.. An exploratory RNA-seq comparison of one matched tumor-adjacent tissue pair confirmed higher tumor abundance of *MALAT1* and the epithelial-to-mesenchymal transition-marker *VIM*, together with lower abundance of cell-adhesion gene *PCDH10*. To investigate the impact of *MALAT1* on thyroid cancer and cellular metabolism, we targeted *MALAT1* in the papillary thyroid cancer cell line TPC1. *MALAT1* knock-down reduced proliferation and migration while enhancing mitochondrial respiration with no changes in glycolysis. Notably, although MALAT1 was not localized within mitochondria, its silencing modulated the expression of transcripts associated with mitochondrial dynamics and mitophagy. Consistent with these results, transcriptomic correlation analysis in the TCGA-THCA cohort showed that MALAT1 expression was largely uncoupled from oxidative phosphorylation and glycolysis gene programs, while negatively correlating with core regulators of mitophagy and mitochondrial dynamics, pointing to a link with mitochondrial quality control rather than direct bioenergetic reprogramming. Our findings highlight *MALAT1* as a contributor to thyroid cancer aggressiveness and reveal a link between *MALAT1* and mitochondrial quality control independent of direct mitochondrial localization. Besides, our results support a tissue-specific mechanism and population-specific role of *MALAT1* in cancer biology.

## Introduction

Thyroid cancer is the most common endocrine malignancy, with approximately 800,000 cases reported worldwide, 8% of which occur in Latin America (1). In Chile, the adjusted incidence of thyroid cancer ranges from 48.7 to 60.8 cases per 100,000 individuals (2), making it the second highest globally after South Korea (2, 3). While multiple subtypes of thyroid cancer have been categorized (4), the well-differentiated forms are the most prevalent (5). These include papillary thyroid cancer, which accounts for 84% of cases, followed by follicular (4%) and oncocytic (2%) subtypes (5).

The incidence of thyroid cancer in Chile has steadily increased over the past decades, a trend also observed worldwide (2). Advances in diagnostic techniques and treatment strategies have led to earlier detection and generally favorable prognoses; however, most patients undergo total thyroidectomy or thyroid lobectomy, procedures associated with high costs (6, 7). In cases where initial evaluations yield indeterminate results, clinical guidelines recommend complementary studies to determine if surgery is needed, including molecular marker analysis (7). While genetic classifiers are currently available (reviewed in (8)), ongoing research aims to identify novel biomarkers to enhance diagnosis and patient management (9, 10).

Several studies have proposed non-coding RNAs (ncRNAs) as potential biomarkers for thyroid cancer (8, 11, 12). ncRNAs are transcripts that are not translated into proteins but function in their native form. This family includes short ncRNAs (e.g., microRNAs, < 200 nucleotides) and long ncRNAs (lncRNAs, >200 nucleotides), both of which play diverse biological roles, including gene regulation in cancer cells (13, 14).

One of the most studied long non-coding RNAs (lncRNAs) in cancer is the Metastasis-Associated Lung Adenocarcinoma Transcript 1 (*MALAT1*). Initially identified as an overexpressed transcript in lung cancer patients (15), *MALAT1* has since been implicated in various malignancies (16). Its role in cancer progression is mediated through multiple mechanisms, including miRNA sponging and interactions with the epigenetic machinery to regulate gene expression (17, 18).

A connection between *MALAT1* and thyroid cancer has been identified, raising questions about its potential as a reliable biomarker for diagnosis and prognosis, as well as its role in cancer progression. *MALAT1* has been quantified in samples from American (19) and Chinese individuals (20–23), showing higher levels in tumor tissues compared to controls; however, no studies have been done on Hispanic population (6).

Regarding its molecular functions, *MALAT1* has been linked to proliferation and migration in thyroid cancer cells through regulation of several genes (20–22), but emerging evidence suggests its involvement in metabolic regulation. Dysregulated cellular energetics is a well-established hallmark of malignancy, as reprogramming of intermediary and energy metabolism is essential for tumorigenesis, proliferation, and invasive capacity (24). In hepatocellular carcinoma, *MALAT1* has been shown to shuttle into mitochondria (25, 26), raising the question of whether the transcript also displays this property in thyroid cancer.

In this study, we analyzed *MALAT1* levels in a cohort of Chilean patient samples and identified a positive correlation with malignancy-associated gene expression and lymphatic infiltration, which can be a sign of a more aggressive disease. Additionally, we report that *MALAT1* upregulation in thyroid cancer is linked to increase proliferation and migration as well as impaired mitochondrial quality control, without direct mitochondrial localization, in the papillary thyroid cancer cell line TPC1.

## Materials and Methods

### Thyroid tissue samples

This study was approved by the Ethics Committee of Universidad Andres Bello and Clinica Indisa. All experiments were performed in accordance with relevant guidelines and regulations. Written informed consent was obtained from all patients prior to sample collection. Patients diagnosed with thyroid cancer from INDISA clinic in Santiago, Chile underwent total thyroidectomy, and tumor and adjacent tissue samples were collected and preserved in RNAlater (Thermo Fisher Scientific, USA) or stored in TRIzol (Thermo Fisher Scientific) at -20°C. Clinical and demographic data were also obtained for each patient.

### RNA extraction

Total RNA was extracted from tissue and cell samples stored in TRIzol using the Direct-zol RNA kit (Zymo Research, USA), according to the manufacturer’s instructions. RNA concentration was measured using a NanoDrop spectrophotometer (Thermo Fisher Scientific), and the 260/280 ratio was recorded. RNA integrity was assessed using a Fragment Analyzer system (Agilent, USA).

### Cell fractionation

To prepare RNA extracts from cytoplasmic and nuclear fractions we followed a modified Dignam *et al* protocol (27). Cells were resuspended in Buffer A at a volume equivalent to 5 pellet volumes. Buffer A consisted of 10 mM Tris (pH 7.9), 1.5 mM MgCl_2_, 10 mM KCl, 0.5 mM DTT, protease inhibitor cocktail, and a 1:100 dilution of RNase inhibitor. The cell suspension was incubated for 10 min at 4°C without agitation. Subsequently, the sample was centrifuged at 2,000 *g* for 3 min, the supernatant was discarded, and Buffer A was added at a volume equal to 2 initial pellet volumes. Cells were mechanically homogenized 10 to 20 times using a syringe equipped with a 20G needle and immediately placed on ice. To separate the cytosolic extract from intact nuclei, the resulting homogenate was centrifuged at 4,000 *g* for 5 min at 4°C. The supernatant (cytosolic extract), while the nuclear pellet was maintained at 4°C. One volume of Buffer B (0.3 M Tris pH 7.9, 0.03 M MgCl_2_, and 1.4 M KCl) was added to the cytosolic supernatant. The mixture was then centrifuged at 15,000 *g* for 30 min at 4°C to clear debris, and the clarified cytosolic supernatant was recovered and RNA was extracted with Trizol. To obtain the nuclear extract, the nuclear pellet was resuspended in 1x RIPA buffer supplemented with protease and RNase inhibitors, incubated on ice for 20 min, and sonicated for 20 s at 20% amplitude using a Qsonica Q800R3 sonicator. Finally, the nuclear lysate was centrifuged at 15,000 *g* for 30 min at 4°C, and the resulting supernatant containing the nuclear extracts was collected and RNA was extracted with Trizol.

### Quantitative real-time PCR

1 µg of total RNA was reverse-transcribed using the SuperScript IV enzyme in the presence of random hexamers (all reagents from Thermo Fisher Scientific). cDNA volumes were adjusted to 100 µL in all reactions. For real-time PCR, cDNA was amplified using the primers shown in **Table 1** with the Brilliant II SYBR Green QPCR Master Mix (Agilent). Relative gene expression levels were calculated using the 2^(-ΔΔCt) method.

**Table 1.**
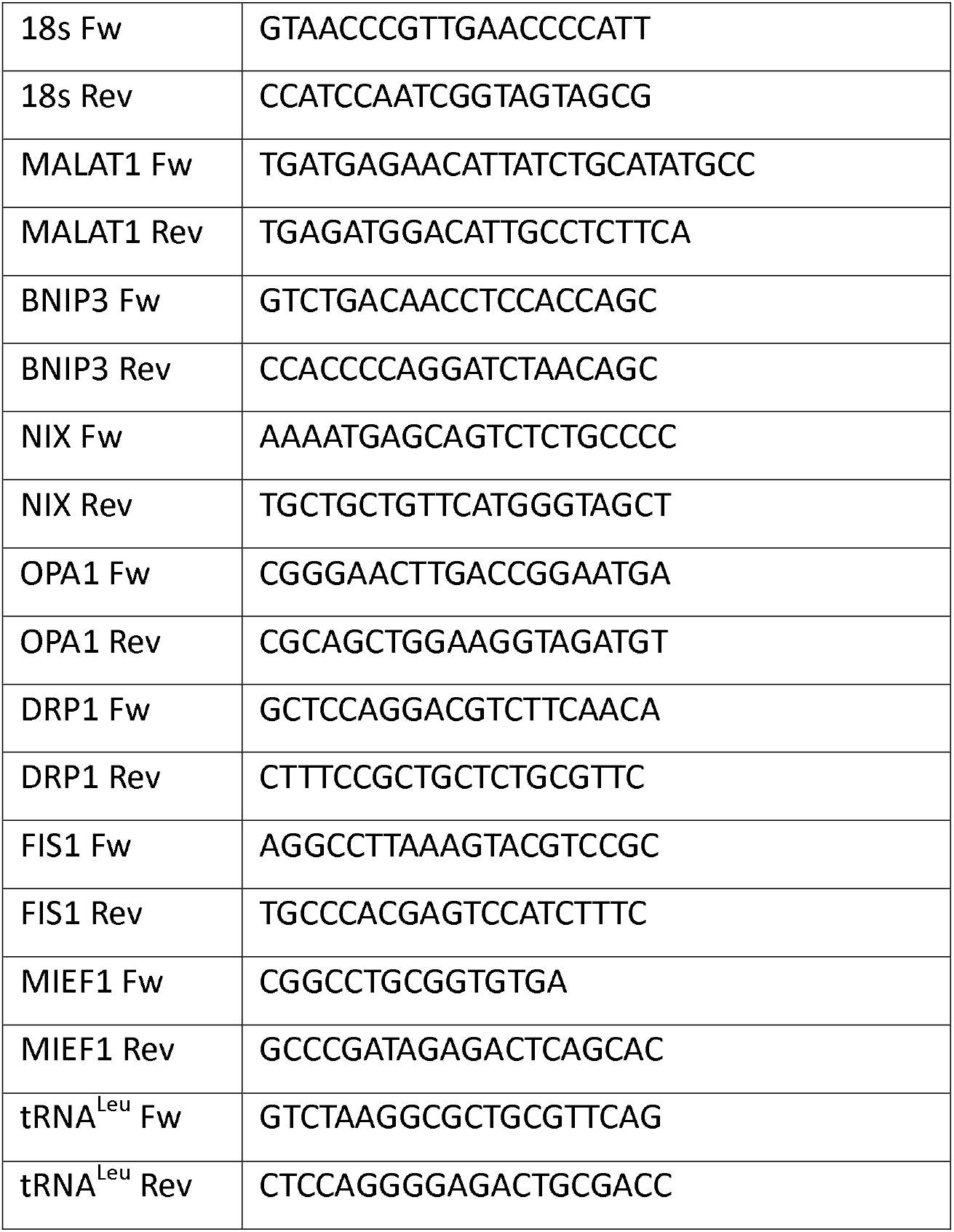
Primers used in this study.

#### RNA sequencing and transcriptome analysis

The RNA sequencing library from one patient was prepared using the MGIEasy RNA Library Prep Kit V3.1 (MGI, China) following rRNA depletion with the Ribo-off rRNA Depletion Kit (Human/Mouse/Rat) (Vazyme, China). Library circularization was performed using MGIEasy Circularization Module V2.0 (MGI). Sequencing was carried out on a DNBSEQ-G400 platform (MGI) using the DNBSEQ-G400RS High-throughput Sequencing Set with single-end 100 bp reads. Sequencing quality was assessed using FastQC (v0.12.0) (28). Adapters and low-quality reads were removed using Trimmomatic (v0.38) (29). Trimmed reads were aligned to the human GRCh38 reference genome using STAR (v2.7.11b) (30). Gene-level raw counts were generated from primary alignments using featureCounts (v2.1.1) (31), assigning reads overlapping annotated exons to gene identifiers in an unstranded single-end configuration. Multimapping and ambiguously assigned reads were excluded. Raw gene counts were normalized using the median-of-ratios method implemented in DESeq2 (v1.52.0). TMM normalization implemented in edgeR (v4.10.1) was additionally used as an independent sensitivity analysis. Descriptive tumor-to-adjacent log2 fold changes were calculated from DESeq2-normalized counts using a pseudocount of 0.5 to avoid undefined ratios for genes with zero counts in one sample. To minimize fold-change estimates driven by very low abundance, downstream analyses were restricted to genes reaching at least 1 TMM-CPM in both samples. Genes with a descriptive log2 fold change calculated from DESeq2-normalized counts ≥1 were classified as having higher normalized abundance in the tumor, whereas genes with a log2 fold change ≤−1 were classified as having lower normalized abundance.

Because only one matched tumor–adjacent tissue pair was sequenced, biological dispersion could not be estimated reliably. Therefore, no gene-level hypothesis tests, p-values, or false-discovery rates were calculated, and the RNA-seq analysis was treated as an exploratory within-patient comparison.

Gene Ontology Biological Process over-representation analysis was performed separately for genes with higher and lower normalized abundance, using genes reaching at least 1 TMM-CPM in both tissues as the analysis universe. After gene-identifier mapping, 11,787 genes from the expressed background, 3,606 genes with higher normalized abundance, and 1,621 genes with lower normalized abundance were represented in the GO analysis. Enrichment was evaluated using a one-sided hypergeometric test and Benjamini–Hochberg correction for multiple testing. Terms with adjusted p-values <0.05 were considered significantly enriched.

### Public Dataset Retrieval and TCGA-THCA Bioinformatic Analysis

To evaluate the scale and generalizability of our findings in an independent cohort, RNA-sequencing expression data (log2 RSEM-normalized) and matched clinical metadata from the TCGA Thyroid Carcinoma project (TCGA-THCA, n ≈ 500) were downloaded from cBioPortal (cbioportal.org). Clinical variables, specifically pathologic lymph node status (N0 vs. N1), were extracted. Gene expression levels for *MALAT1* and three candidate targets (CDH2, VIM, and PCDH10) were parsed across both the unstratified whole cohort (n ≈ 500) and the node-positive clinical subset (N1 stage, n = 257). Bivariate associations were assessed using Spearman rank correlation coefficients (ρ), with raw *p-*values adjusted for multiple testing using the Benjamini-Hochberg false discovery rate (*padj*). All bioinformatic analyses and statistical visualizations were conducted in R (v4.3.0) using *tidyverse* and *ggpubr* packages. To further evaluate whether *MALAT1* expression is associated with lymphatic infiltration at a population scale, we performed a multiple linear regression analysis using the TCGA-THCA cohort described above, with *MALAT1* expression as the outcome and lymph node status (N1 vs. N0), age, gender, and TCGA-THCA molecular subtype as predictors. Additionally, to explore associations between *MALAT1* and mitochondrial quality-control pathways, we computed Pearson correlation matrices between *MALAT1* expression and curated gene sets representing oxidative phosphorylation (26 genes), glycolysis (23 genes), and mitochondrial dynamics/mitophagy (30 genes), stratified by pathologic lymph node status (N0, n = 226; N1, n = 222); correlation coefficients and raw *p*-values are reported for each gene pair.

### Cell culture and transfection of Antisense Oligonucleotides

The human papillary thyroid carcinoma cell line TPC1 (Sigma-Aldrich, USA) was used in this study(32). The TPC1 cell line was originally sourced from the laboratory of Dr. J. A. Fagin (Memorial Sloan Kettering Cancer Center, New York, NY) and generously provided by Dr. Hernán González (Pontificia Universidad Católica de Chile). Cell line authenticity and genetic identity were verified prior to use following established benchmarks(33). Mycoplasma contamination was discarded periodically using the Mycoplasma PCR Detection Kit (Applied Biological Materials Inc., Canada). Cells were cultured in RPMI 1640 HG medium (Capricorn Scientific, Germany) supplemented with 10% fetal bovine serum (Gibco, USA), 10 mM pyruvate (Capricorn Scientific), and 100 U/mL penicillin-streptomycin (Gibco). The cultures were maintained at 37°C in a humidified incubator with 5% CO₂. *MALAT1* was silenced in TPC1 cells for 72 hours using commercial LNA-GapmeRs antisense oligonucleotides (ASOs), catalog number: LG00000003-DDB (Qiagen, USA), which were transfected at a concentration of 10 nM using Lipofectamine 3000 (Thermo Fisher Scientific), following the manufacturer’s instructions. A commercial scrambled sequence ASO, catalog number: LG00000002-DDB (Qiagen), was used as a control. Silencing of *MALAT1* was confirmed by RT-qPCR as described above. Since the ASOs were conjugated to the 5’-FAM fluorescent dye, transfected cells were imaged using a Nikon Eclipse epifluorescence microscope (Japan).

### Cell proliferation study

A total of 100,000 cells were seeded per well. Every 24 hours, cells were counted using a Neubauer hemocytometer (Paul Marienfeld, UK) under an optical microscope in the presence of 0.4% trypan blue solution (Invitrogen).

### Cell cycle analysis

Cells were incubated with the DNA intercalating agent Propidium Iodide (Sigma-Aldrich) at a concentration of 40 µg/mL. To preserve the fluorescent ASO within the cells, the cells were fixed with 1% paraformaldehyde (Sigma-Aldrich). Cells were then analyzed by flow cytometry using a FACSAria III cytometer (BD Biosciences, USA) with an excitation wavelength of 488 nm and an emission wavelength of 625 nm. The proportion of cells in the G0/G1, S, or G2/M phases was determined from 10,000 events.

### Cell migration study

Cells were seeded onto inserts in Transwell® plates (Corning, USA). At 4-hour intervals, cells that had passed through the porous membrane were fixed and stained with crystal violet (Merck-Millipore, USA). Images were captured using a BioTek Lionheart FX Automated Microscope (Agilent Technologies).

### TMRE analysis

Mitochondrial membrane potential was assessed by staining cells with 10 nM TMRE (Thermo Fisher Scientific) in a non-quenching mode for 30 minutes at 37°C. Cell imaging was performed using a confocal Olympus Fluoview 1000 microscope. Single confocal images were acquired at 60x magnification. Images were analyzed using Fiji software, and the mean fluorescence intensity of the entire cell was measured to determine mitochondrial membrane potential.

### Metabolic analysis (ECAR, OCR)

Extracellular acidification rates (ECAR) and Oxygen consumption (OCR) were measured using the Seahorse XF Pro Extracellular Flux Analyzer (Agilent), as described in (34). Cells were plated or 20,000 (XF Pro) cells per well in XF Pro M plates (XF Pro). Basal respiration, ATP-linked respiration, proton leakage, maximal respiration, and non-mitochondrial respiration were measured by the sequential addition of 5 mM glucose, 1 μM oligomycin, 0.75 μM FCCP, and 1 μM rotenone/1 μM antimycin A.

### RNA Fluorescence in-situ hybridization (RNA-FISH) and mitochondrial detection

Live cells were treated with MitoTracker® Green dye (Thermo Fisher Scientific) for 30 minutes at 37°C before fixing cells with 3.7% formaldehyde (Sigma-Aldrich) and permeabilizing with 70% ethanol (Merck) for RNA-FISH protocol. For *MALAT1* detection, we utilized probes from Biosearch Technologies (Stellaris® FISH Probes, Human *MALAT1* with Quasar® 570 Dye, Cat. SMF-2035-1, USA), comprising a mix of oligonucleotides targeting the entire RNA molecule. Hybridization steps with the specific probes were performed according to the Stellaris RNA FISH protocol. Hybridization was conducted at 37Ö°C for 16 hours; washes were performed with specific buffers to remove unbound probes; and nuclei were stained with NucBlue™ Fixed Cell ReadyProbes™ Reagent (DAPI, Thermo Fisher Scientific) before mounting. Images were captured using a confocal Olympus Fluoview 1000 microscope.

### Western blot analysis

Cell pellets were homogenized in NP-40 lysis buffer containing 150 mM NaCl, 1% NP-40, 50 mM Tris (pH 8.0), 0.1% SDS, along with a protease inhibitor cocktail (Sigma-Aldrich) and PMSF (Thermo Fisher Scientific). Total proteins were separated in a 15% polyacrylamide gel containing 2,2,2-trichloroethanol, which enables fluorescent visualization of proteins under UV light exposure,s serving as a protein loading control (35). Blots were blocked for 1 hour in 5% non-fat dry milk in TBS-T buffer (Tris-buffered saline, pH 7.4, 0.2% Tween-20) during 1 hr, and then incubated with the primary antibodies (mouse anti total OXPHOS antibody cocktail, Abcam, USA, # ab110413; rabbit anti GAPDH, Cell signaling, USA, #2118) in TBS-T overnight at 4 °C. After three washes with TBS-T for 5 min each, blots were incubated with the HRP-conjugated secondary antibody (goat anti Mouse IgG (H+L) HrP, Invitrogen #31432; goat anti Rabbit IgG (H+L) HrP, Invitrogen # 31460) in blocking solution for 1 hour. Finally, membranes were treated with SuperSignal™ West Femto substrate (Thermo Fisher Scientific) and visualized using the Alliance Q9 Advanced UVITEC imaging system. Band densitometric analysis was performed using the NINE ALLIANCE UVITEC software.

### Statistical analysis

All calculations, data analyses, statistical tests (Paired Student’s t-test, Mann-Whitney or Kruskal-Wallis as needed), and plots were conducted using GraphPad Prism® version 10, Jamovi or R. To compare paired *MALAT1* fold-change values between tumor and adjacent non-tumor tissue, we used a paired t-test and Gardner-Altman estimation plot. To identify independent clinical predictors of *MALAT1* expression (ΔΔCt), a multiple linear regression model was adjusted for age, tumor size, sex, ATA risk category, and lymphatic infiltration status as predictors. Similarly, the relationship between TMRE mean fluorescence intensity and transfection condition (control ASO vs. anti-*MALAT1* ASO), adjusted for GFP fluorescence as a covariate, was assessed by linear regression in Jamovi.

## Results

### MALAT1 is highly expressed in Chilean patients with lymphatic infiltration

We recruited 16 Chilean patients who underwent surgery following a thyroid cancer diagnosis. As summarized in **Table 2**, eleven patients were women, in line with the higher prevalence of thyroid cancer in Chilean females (2). One patient had a benign tumor, while fourteen were diagnosed with papillary thyroid carcinoma (including the aggressive trabecular and tall cell subtypes) the most common histological variant in Chile (2). Patients exhibited different risk levels according to the American Thyroid Association (ATA). Twelve tumors were larger than 20 mm, seven exhibited lymphatic infiltration, and ten were positive for lymph node metastasis (**Table 2, Supplementary Table S1**).

**Table 2.** Frequency data of thyroid cancer patients enrolled in this study.

| Variable | Number of Patients |
| --- | --- |
| **Type of cancer** |  |
| *Benign\** | 1 |
| Papillary | 12 |
| Follicular subtype | 1 |
| Tall cells subtype | 1 |
| Trabecular | 1 |
| **Age** |  |
| < 50 | 10 |
| >50 | 5 |
| **Gender** |  |
| Male | 4 |
| Female | 11 |
| **Primary tumour** |  |
| T1a/b | 3 |
| T2 | 10 |
| T3 | 1 |
| T4a | 1 |
| **ATA\*\* risk** |  |
| Low (1) | 4 |
| Intermediate (2) | 9 |
| High (3) | 2 |
| **Tumour size** |  |
| < 20 mm | 3 |
| ≥ 20 mm | 12 |
| **Lymphatic infiltration** |  |
| Negative | 8 |
| Positive | 7 |
| **Lymph node metastases** |  |
| Negative | 5 |
| Positive | 10 |

\*The benign-tumor patient is excluded from tumor-staging-related rows.
\*\* ATA: American Thyroid Association.

To determine whether our cohort exhibited elevated *MALAT1* lncRNA levels, as previously reported in thyroid cancer patients from American (19) and Chinese populations (20–23), we performed quantitative RT-PCR analysis, normalizing *MALAT1* levels to adjacent non-tumor tissue from the same patient. As shown in the Gardner-Altman plot (**Fig. 1a**) and **Supplementary Table S1**, *MALAT1* expression relative to non-tumoral paired tissue varied among patients. When PCR data was analyzed by a paired t-test (mean difference between adjacent and tumoral tissues) no significant differences were found (**Fig. 1a**).

**Figure 1.**
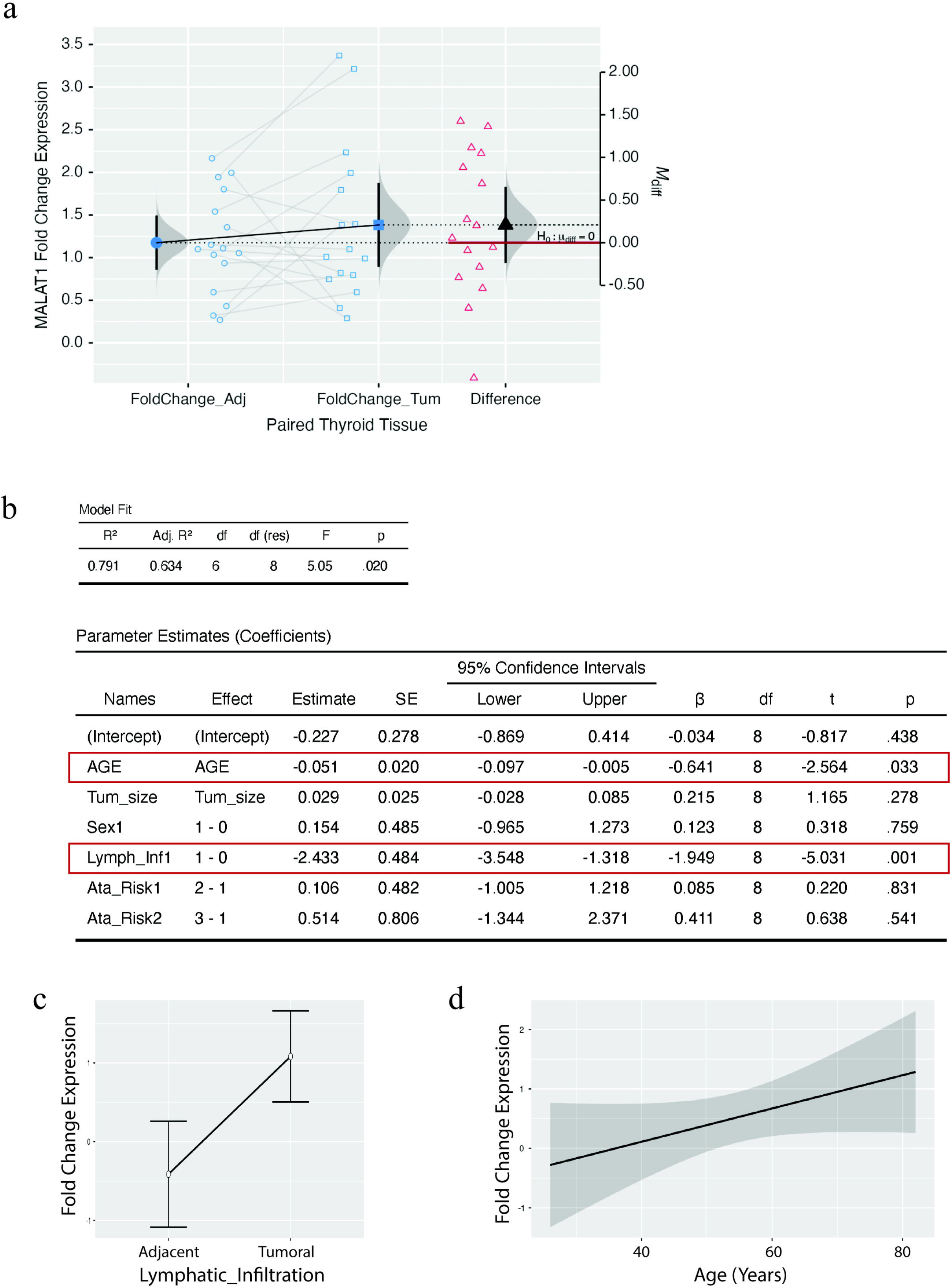
MALAT1 gene expression in thyroid cancer patient samples. Tumor and adjacent non-tumor thyroid tissues were collected from patients diagnosed with thyroid cancer. Total RNA was extracted, reverse transcribed to cDNA and analyzed by quantitative PCR using *MALAT1*-specific primers. GAPDH served as housekeeping. **(a)** Paired comparison of *MALAT1* fold-change expression between adjacent non-tumor and tumor tissue, displayed as a Gardner-Altman estimation plot; individual paired values are connected by lines, and the mean difference with its 95% confidence interval is shown on the right axis. A statistical paired t-test was performed and no significant differences were found in MALAT1 expression between adjacent and non-tumoral tissue (p>0.05). **(b)** Multiple linear regression model of MALAT1 ΔΔCt values, with age, tumor size, sex, ATA risk category, and lymphatic infiltration status as predictors. Model fit statistics and parameter estimates (unstandardized and standardized coefficients, 95% confidence intervals, and p-values) are shown; because ΔΔCt is inversely related to relative expression, negative coefficients correspond to higher MALAT1 expression. **(c)** Model-estimated *MALAT1* fold-change expression by lymphatic infiltration status, derived from the multiple regression model shown in (b). **(d)** *MALAT1* fold-change expression as a function of patient age, with a fitted linear regression line and 95% confidence band, likewise derived from the model in (b).

To identify independent clinical predictors of *MALAT1* expression, we fitted a multiple linear regression model using *MALAT1* ΔΔCt difference values between adjacent and tumoral tissue as the dependent variable, with age, tumor size, sex, ATA risk category, and lymphatic infiltration status as predictors (**Fig. 1b-d**). The overall model was statistically significant (R² = 0.791, adjusted R² = 0.634, p = .020) meaning that 63,4% of *MALAT1* variability is explained by the model. Because relative *MALAT1* expression is inversely related to ΔΔCt (fold change = 2 raised to the power of −ΔΔCt), a negative regression coefficient corresponds to higher *MALAT1* expression. Lymphatic infiltration was the strongest independent predictor (β = -1.949, p = .001), confirming higher *MALAT1* expression in tumors with lymphatic infiltration after adjusting for age, tumor size, sex, and ATA risk (**Fig. 1c**). Age was independently associated with ΔΔCt (β = -0.641, p = .033), indicating higher *MALAT1* expression in older patients, consistent with the positive trend between fold-change expression and age (**Fig. 1d**). Tumor size, sex, and ATA risk category did not reach statistical significance (all p > .05).

To directly test whether the relationship between *MALAT1* expression and lymphatic infiltration observed in our Chilean cohort generalizes to a larger population, we modeled *MALAT1* expression itself as the outcome in the TCGA-THCA cohort, mirroring the multiple regression approach used in our own cohort (**Fig. 1b**), with lymph node status, age, gender, and molecular subtype (BRAF-like as reference, compared with RAS-like and Other/Not Applicable) as predictors, restricted to patients with complete covariate data (N = 419) (**Supplementary Figure S1**). The overall model explained a modest proportion of variance (R = 0.254, R² = 0.065). Lymph node status was not a significant predictor of *MALAT1* expression in this larger, unselected cohort (estimate = 0.099, p = .266), nor were age (estimate = -0.003, p = .280) or gender (estimate = 0.062, p = .506). Molecular subtype was the only strong predictor, with the Braf-like (estimate -0,4, p<0.01) subtype associated with lower MALAT expression relative to Ras-like and Other/Not Applicable subtypes. These results indicate that the univariate association between *MALAT1* and lymphatic infiltration observed in our small Chilean cohort does not generalize to a larger, subtype-heterogeneous population, where molecular subtype (rather than nodal status) is the dominant driver of *MALAT1* expression variability.

### Case study: Exploratory normalized transcriptomic profile ofa high-risk (ATA 3) thyroid tumor

To explore the transcriptomic profile associated with a tumor exhibiting high *MALAT1* levels, we analyzed tumor tissue and its matched adjacent non-tumor tissue from patient 16, classified as ATA level 3 (high risk) (**Supplementary Table S1**). Because only one matched pair was sequenced, this analysis was considered descriptive and exploratory.

Following realignment and gene-level quantification, 33,121 annotated gene features were quantified, of which 25,956 had at least one assigned count (**Supplementary Table S2**). Raw counts were normalized using the DESeq2 median-of-ratios method, with TMM normalization used as an independent sensitivity analysis. The two normalization approaches produced highly concordant descriptive fold-change estimates (Spearman’s ρ = 0.99994).

To minimize fold-change estimates driven by very low abundance, descriptive analyses were restricted to genes reaching at least 1 TMM-CPM in both tissues. Under this criterion, 15,372 genes were retained. Of these, 4,407 showed at least a twofold higher normalized abundance and 2,399 showed at least a twofold lower normalized abundance in the tumor, whereas 8,566 showed changes smaller than twofold. These categories represent descriptive within-patient abundance differences and do not constitute statistically significant differential-expression calls. Gene Ontology analysis (**Fig. 2a**) identified DNA damage tolerance as the only significantly enriched selected cancer-related Biological Process among genes with higher normalized abundance in the tumor (23 genes; adjusted p = 0.0425). Genes with lower normalized abundance were enriched for humoral immune response (37 genes; adjusted p = 8.43 × 10⁻^9^), cellular component assembly involved in morphogenesis (27 genes; adjusted p = 4.81 × 10⁻⁵), G protein-coupled receptor signaling pathway (98 genes; adjusted p = 9.76 × 10⁻⁵), extracellular matrix organization (52 genes; adjusted p = 6.09 × 10⁻⁴), and cilium movement involved in cell motility (27 genes; adjusted p = 0.0103) (**Fig. 2a**). The genes showing the largest positive and negative descriptive log2 fold changes are shown in **Fig. 2b**.

**Figure 2.**
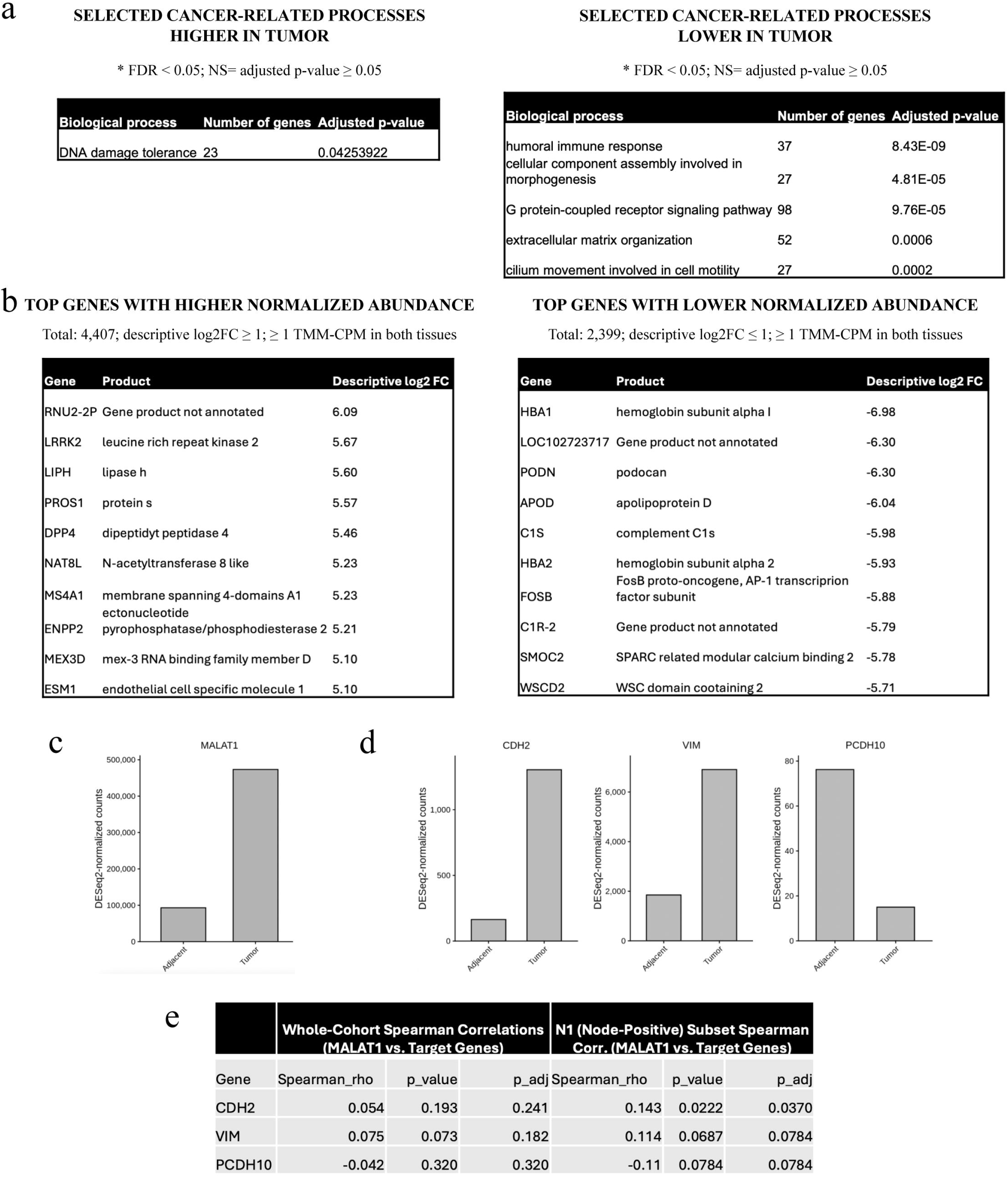
Case study: Transcriptomic comparison of a high-risk (ATA 3) thyroid tumor and matched adjacent tissue. Tumor and matched adjacent non-tumor tissue from one high-risk (ATA 3) patient were analyzed by RNA sequencing. Gene-level counts were normalized using the DESeq2 median-of-ratios method, with TMM normalization used as an independent sensitivity analysis. Descriptive analyses were restricted to genes reaching at least 1 TMM-CPM in both tissues. **(a)** Selected cancer-related Gene Ontology Biological Process terms significantly enriched among genes with higher or lower normalized abundance in the tumor. Pathway-level p-values were adjusted using the Benjamini–Hochberg method. **(b)** Genes with the highest positive or negative descriptive log2 fold change calculated from DESeq2-normalized counts. **(c)** DESeq2-normalized MALAT1 counts in the matched adjacent and tumor tissues. **(d)** DESeq2-normalized counts for selected EMT-associated genes (*CDH2*, *VIM*, and *PCDH10*) and the cell-adhesion gene PCDH10. Because only one matched pair was sequenced, no gene-level p-values orfalse-discovery rates were calculated, and all gene-level comparisons are descriptive. **(e)** Gene expression levels for *MALAT1* and the three genes analyzed in (d) (*CDH2*, *VIM*, and *PCDH10*) were parsed across both the unstratified whole cohort and the node-positive clinical subset (N1). Bivariate associations were assessed using Spearman rank correlation coefficients (ρ), with raw p-values adjusted for multiple testing using the Benjamini-Hochberg false discovery rate (p-adj).

We then focused on selected genes of interest. *MALAT1*, which was more abundant in this tumor relative to the matched adjacent tissue by RT-qPCR (**Fig. 1a, Supplementary Table S1**), also displayed higher normalized abundance in the RNA-seq analysis. DESeq2 normalization yielded a descriptive log2 fold change of 2.35, corresponding to a 5.08-fold higher abundance in the tumor. TMM normalization produced a closely comparable estimate of 5.19-fold (**Fig. 2c, Supplementary Table S2**). Selected EMT-associated genes also showed higher normalized abundance in the tumor, including CDH2 (N-cadherin; descriptive log2FC = 2.99), and VIM (Vimentin; log2FC = 1.90). Conversely, PCDH10 (Protocadherin 10), a cell-adhesion gene, showed lower normalized abundance (log2FC = −2.31) (**Figure 2d, Supplementary Table S2**). These observations represent a within-patient transcriptomic profile and require validation in additional biological samples.

To determine whether the co-expression relationships between *MALAT1* and EMT/adhesion markers observed in our clinical cohort generalize to population-scale datasets, we analyzed RNA-seq data from the TCGA-THCA cohort. In the unstratified whole cohort, rank correlations between *MALAT1* and individual targets were generally weak or non-significant (**Fig. 2e**, “Whole cohort”). Because unstratified bulk sequencing aggregates indolent, non-metastatic neoplasms alongside aggressive variants (potentially diluting invasion-specific co-expression networks) we next restricted our analysis to tumors with documented regional lymph node metastasis (N1 stage, n=257). Within this metastatic subset, filtering out indolent background cases revealed distinct co-expression dynamics. Notably, *MALAT1* demonstrated a statistically significant positive correlation with the markers (**Fig. 2e**, “N1 Subset”).

### MALAT1 modulates proliferation and migration in the papillary thyroid cancer cell line TPC1

Given the observed *correlation* between *MALAT1* levels and lymphatic infiltration (**Fig. 1 b-c**), as well as its elevated levels in a patient with metastasis, we sought to determine whether *MALAT1* contributes to the aggressive properties of thyroid cancer cells. We performed antisense oligonucleotide (ASO)-mediated silencing of the *MALAT1* RNA in the papillary thyroid cancer cell line TPC1, using LNA-GapmeRs conjugated to the green fluorescent dye 6-FAM.

Cellular uptake of *MALAT1* ASOs and their accumulation in the nucleus were confirmed by fluorescence microscopy (**Fig. 3a**). The negative control ASOs were distributed homogeneously within the cells (**Fig. 3a, left**), while the fluorescence of anti-*MALAT1* ASOs was concentrated in foci within the nucleus known as “speckles” (**Fig. 3a, right**), a well-known pattern for this transcript (more below) (36). Additionally, we determined that a concentration of 10 nM ASO was the minimal amount required for maximal *MALAT1* silencing in TPC1 cells, determined by quantitative RT-PCR (∼85–90% knock-down relative to control ASO, **Fig. 3a, far right**).

**Figure 3.**
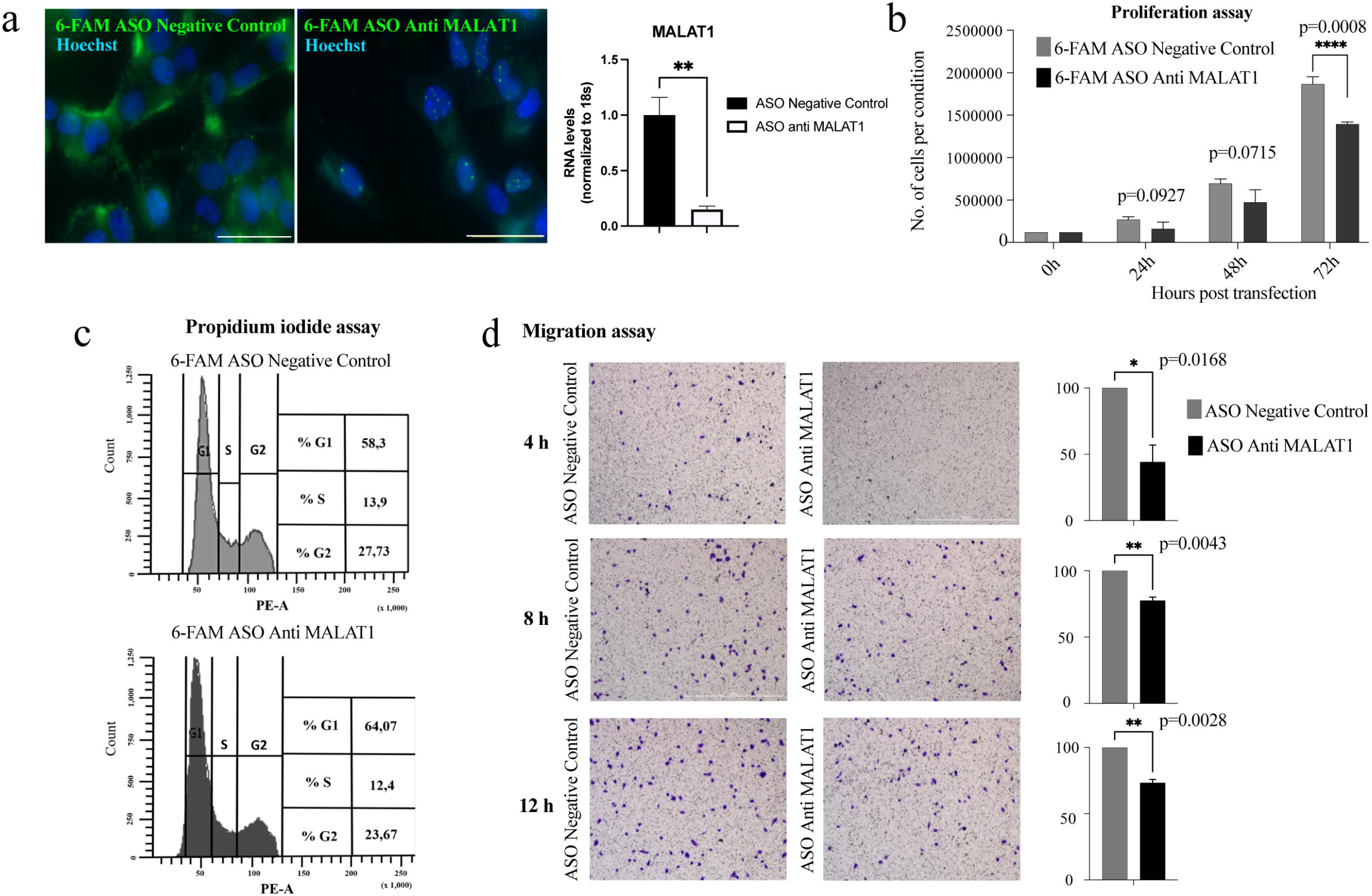
*MALAT1* modulates proliferation and migration in the papillary thyroid cancer cell line TPC1. *MALAT1* was silenced in TPC1 cells using specific antisense oligonucleotides (ASOs) conjugated to the green fluorescent dye 6-FAM. A commercial ASO negative control was included. (a) Left: ASOs uptake was confirmed by epifluorescence microscopy. Images were captured 48 h post-transfection. Nuclei were counterstained with Hoechst (scale bar = 15 μm. A representative result from at least 6 biological replicates, each analyzed in triplicate, is shown). **Right:** *MALAT1* levels were quantified by RT-qPCR using specific primers. 18s served as housekeeping. **(b)** Proliferation assay. Cells were counted every 24h post-transfection. Bars represent the mean ± S.D (n = 3 biological replicates, p<0.01, Student’s t-test). **(c)** Cell cycle analyses. Cells were fixed with 1% paraformaldehyde, stained with 40 μg/mL Propidium Iodide and subjected to cell cytometry, counting 10,000 events. A representative result from 3 biological replicates is shown. **(d)** Migration assays. Those were performed using Transwell chambers, with cells incubated with the ASO for 48h, seeded onto Transwell membranes and then collected at 4, 8, and 12 h. Images show a representative field from the full membrane. Bars represent percentage of cells with respect to control (n = 3 biological replicates, p values determined by Student’s t-test).

To evaluate the impact of *MALAT1* silencing on TPC1 cell proliferation, we quantified cell numbers every 24 hours. A reduction in proliferation was apparent at 48 hours and became statistically significant at 72 hours (**Fig. 3b**). Cell cycle analysis using propidium iodide staining revealed that *MALAT1*-silenced cells accumulated in the G1/S phase at 48 hours, accompanied by a decrease in the G2 phase (**Fig. 3c**). Given the association between *MALAT1* levels and lymphatic infiltration, together with the higher normalized abundance of *VIM* in the exploratory RNA-seq profile (**Fig. 2d**), we next examined the effect of *MALAT1* silencing on cell migration. In Transwell assays, cells incubated with ASOs for 48h and then seeded onto Transwell plates showed a significant reduction in migration following *MALAT1* knock-down (**Fig. 3d**; times indicate hours of cells on the membrane).

In summary, our findings indicate that *MALAT1* knock-down is sufficient to reduce proliferation and migration in TPC1 cells.

### MALAT1 expression is associated with decreased mitochondrial respiration and quality control in TPC1 cells via a non-mitochondrial localization

While the contribution of *MALAT1* to proliferation, cell cycle progression, and migration has been documented in thyroid cancer (20–22) and various other cancer types (16), its role in tumor cell metabolism remains largely unexplored, with studies to date primarily focused on hepatocellular carcinoma (25, 26). Given previous evidence linking *MALAT1* to metabolic regulation (37, 38), we assessed whether *MALAT1* disruption affects cellular metabolism in TPC1 cells .

We first conducted a non-quenching TMRE assay to assess mitochondrial membrane potential (**Fig. 4a**). Whole-cell TMRE fluorescence intensity was measured in positively transfected (GFP+) TPC1 cells. The extent of *MALAT1* silencing is dependent on the amount of ASO anti-*MALAT1* uptake by the cells. Therefore, fluorescent intensity data were collected for both the green (ASO) and red (TMRE) signals and analyzed using linear regression. The dependent variable was the TMRE mean fluorescence, while the transfection conditions (both control ASO and ASO anti *MALAT1*) served as the predictor variables, and GFP fluorescence was included as a covariate. Confocal images, histogram data distribution, and box plots clearly demonstrated a reduction in membrane potential under the *MALAT1* knockdown condition (**Fig. 4a**). The regression model indicated that 55.7% (p < 0.01) of the variability in membrane potential could be explained by the ASO anti *MALAT1* condition, as evidenced by the determination coefficient (R²). Additionally, TMRE decreased by 840.472 fluorescent units when adjusted for GFP fluorescence in the silenced *MALAT1* condition, (p < 0.01) as shown by the model coefficients (Estimate).

**Figure 4:**
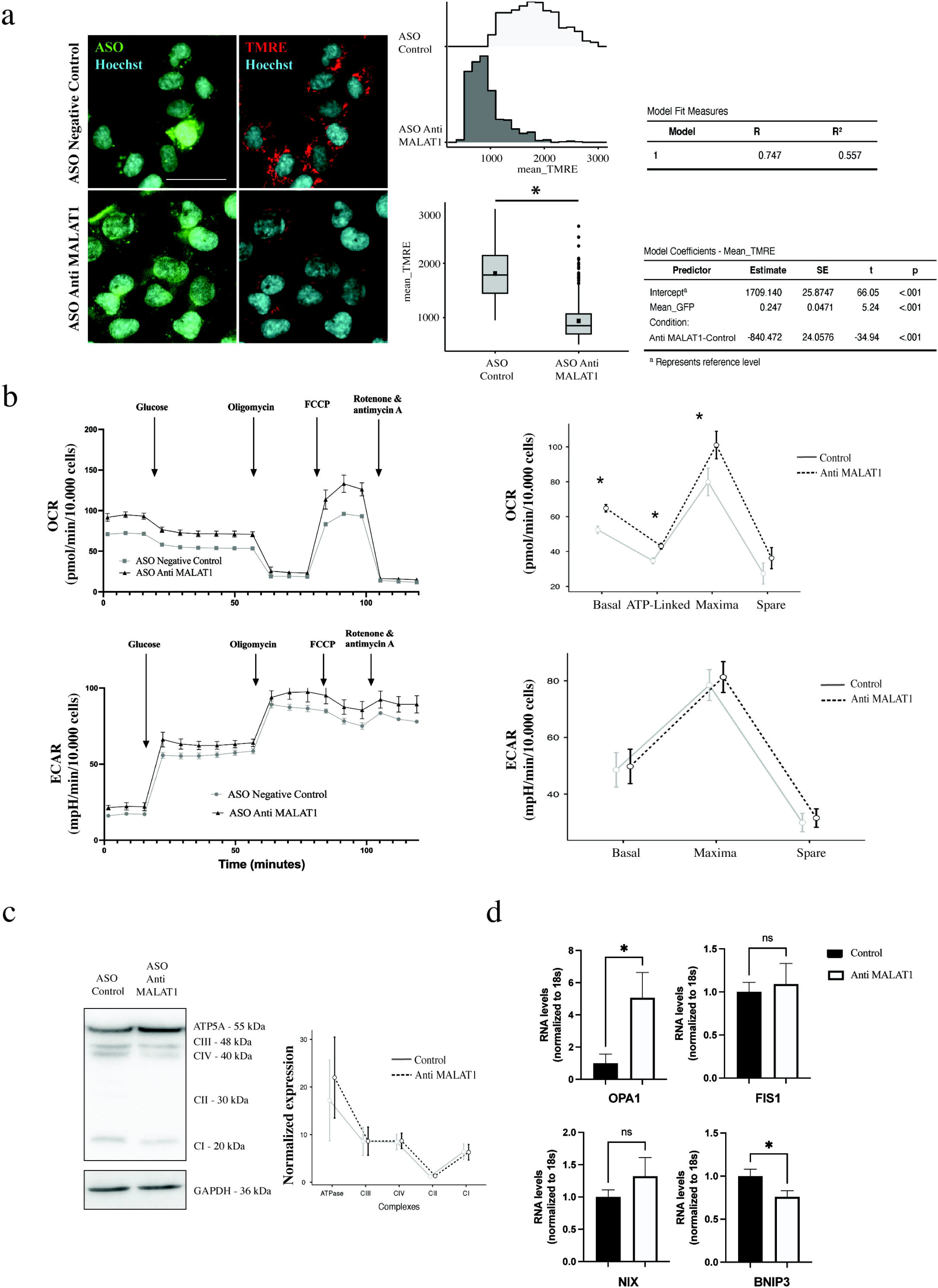
*MALAT1* down-regulation in linked to changes in metabolism of TPC1 cells. *MALAT1* was silenced in TPC1 cells using fluorescent ASOs. The respective negative control is also shown. (a) Mitochondrial membrane potential. It was assessed 72 h post-transfection by staining ASO-containing cells (green) with 10 nM TMRE (red). Nuclei were counterstained with Hoechst (scale bar = 20 μm). *Left:* Images of a representative field. *Center:* Whole-cell TMRE fluorescence intensity was measured in positively transfected TPC1 cells and data were collected for both the green (ASO) and red (TMRE) signals. Histogram data and box plots are shown to depict signal intensity distribution. *Right:* The regression model indicates that 55.7% of the variability in membrane potential could be explained by the ASO anti *MALAT1* condition, as evidenced by the determination coefficient (R²). Also, a model coefficients analysis is shown. **(b)** Oxygen consumption rate (OCR) and Extracellular acidification rate (ECAR) analyses. Representative plots from three biological replicates are shown. **(c)** The expression levels of electron transport chain complexes (C) and the ATP synthase (ATP5A) were assessed by Western blot. One representative blot is shown from three independent experiments. Full-length blots and protein loading controls are shown in **Supplementary Fig. S2**. Densitometric analysis was performed in ImageJ. **(d)** The expression of nuclear-encoded mitochondrial transcripts was determined by RT-qPCR. Bars represent the mean ± S.D (n = 3 biological replicates, Welch’s t-test, *: p<0.05, ns = non-significant differences)

A reduction in membrane potential is typically associated with an increase in ATP synthase activity to meet ATP demand, which subsequently leads to increased oxygen consumption by the electron transport chain (39). To investigate this hypothesis, we assessed the oxygen consumption rate (OCR) and the extracellular acidification rate (ECAR) using Seahorse technology (**Fig. 4b**). As expected, cells with decreased *MALAT1* exhibited a significant increase (p < 0.05) in basal, ATP-linked, and maximal respiration, with no differences observed in spare respiration (**Fig. 4b**). Regarding ECAR, a measure of glycolysis, no changes were noted at basal, maximal, nor spare glycolysis. Furthermore, we measured the protein expression of the respiratory complexes by Western blot. No significant differences were observed in the levels of CI, CII, CIII, CIV and ATP synthase (**Fig. 4c; Supplementary Fig. S2**).

Mitochondrial fitness depends on both OXPHOS and quality control system (dynamics and mitophagy) (40) To investigate this further, we quantified a panel of nuclear-encoded mitochondrial transcripts in *MALAT1* knock-down cells (**Fig. 4d**). *OPA1* transcript, coding for a mitochondrial fusion protein, was significantly increased (*p* = 0.0335) while the mitophagic transcript BNIP3 was significantly decreased ( *p* = 0.0180). The other transcripts (FIS*1*, mitochondrial fission, and *NIX*, mitophagy) did not significantly change (**Fig. 4d**)

To place these bioenergetic findings within a broader molecular context, we examined the relationship between *MALAT1* expression and curated gene sets representing oxidative phosphorylation, glycolysis, and mitochondrial dynamics/mitophagy in the TCGA-THCA cohort, stratified by pathologic lymph node status (**Fig. 5**). Correlations between *MALAT1* and both OXPHOS- (**Fig. 5a**) and glycolysis-related transcripts (**Fig. 5b**) were weak and largely inconsistent in node-negative (N0) tumors, and remained comparatively weak in node-positive (N1) tumors, indicating that *MALAT1* expression is largely decoupled from bulk bioenergetic gene expression at the population level. In contrast, correlations with mitochondrial dynamics and mitophagy transcripts were stronger and predominantly negative in the **N1** subset, particularly for core autophagy/mitophagy effectors (**Fig. 5c**). This pattern is consistent with our functional data implicating *MALAT1* in mitochondrial quality-control processes rather than in direct regulation of OXPHOS or glycolytic gene expression.

**Figure 5.**
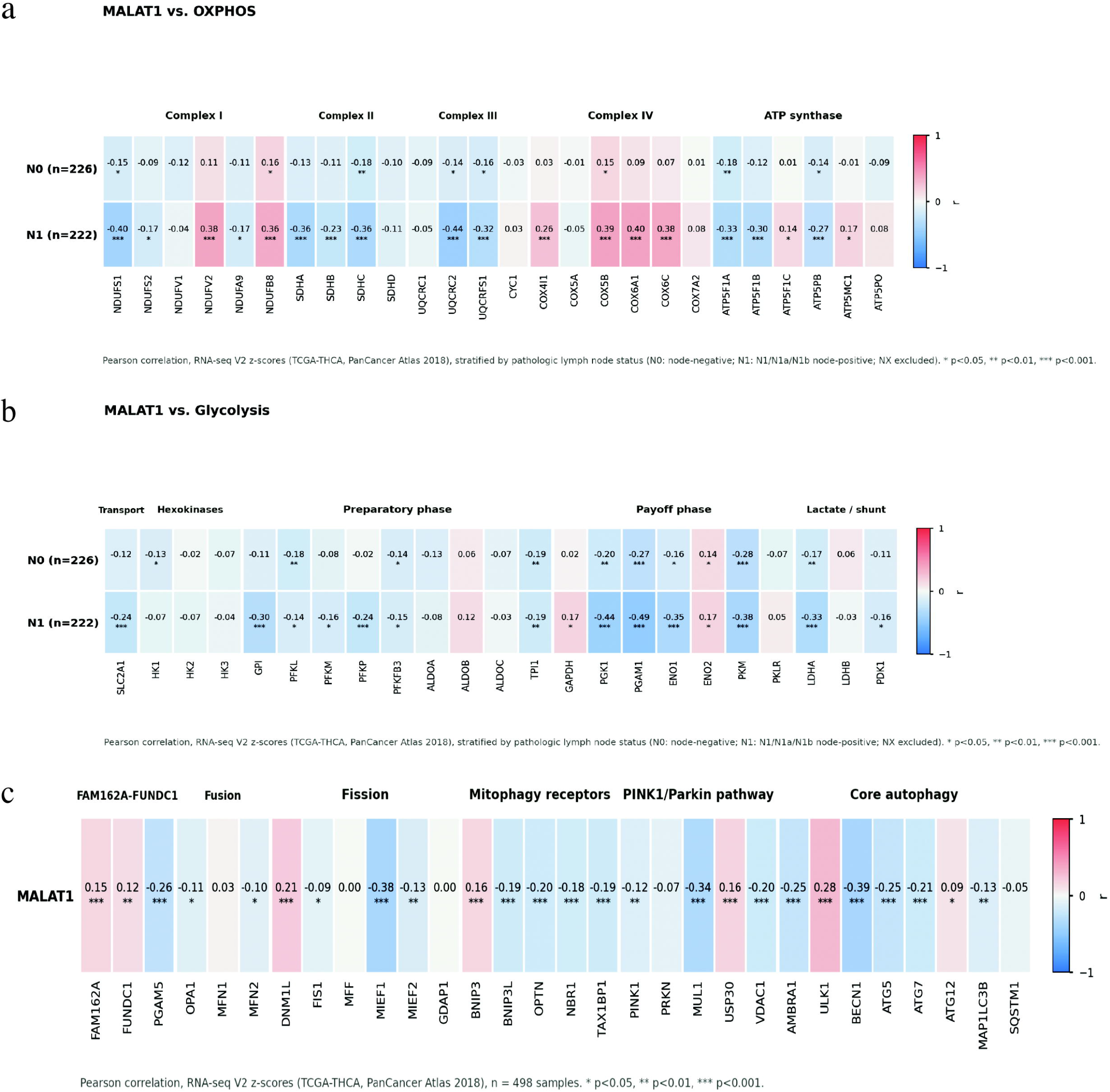
MALAT1 expression is decoupled from bioenergetic gene expression but correlates with mitochondrial dynamics/mitophagy transcripts. Pearson correlation matrices between *MALAT1* expression and curated gene sets representing oxidative phosphorylation, glycolysis, and mitochondrial dynamics/mitophagy in the TCGA-THCA cohort, stratified by pathologic lymph node status (N0, n = 226; N1, n = 222). Asterisks indicate raw p-value significance thresholds for each gene pair.

Although *MALAT1* has previously been reported to shuttle to mitochondria and upregulate mitochondrial metabolism in HepG2 hepatocarcinoma cells (25, 41), we did not observe mitochondrial localization in TPC1 cells. To evaluate its subcellular distribution, we performed RNA-FISH coupled with MitoTracker staining. Confocal microscopy analysis revealed that *MALAT1* was exclusively localized to the nucleus in 90% of TPC1 cells (295 of 329 cells analyzed across four biological replicates), consistent with its canonical nuclear retention (36) (**Fig. 6a, top; Fig. 6b**). An extranuclear MALAT1 signal was detected in 6% of cells (19 cells); however, this signal showed no co-localization with mitochondria (**Fig. 6a, bottom; Fig. 6b**). In the remaining 4% of cells, we did not detect *MALAT1* signal (14 cells). To independently corroborate these imaging results, we performed biochemical cell fractionation followed by RT-qPCR. These experiments confirmed a marked enrichment of *MALAT1* in the nuclear fraction (**Fig. 6c, left**; *p*<0.0001). As a cross-contamination control we used tRNA^Leu^ levels, which are significantly enriched only in the cytoplasmic fraction (**Fig. 6c, right**; *p*<0.0001). Overall, our results demonstrate that *MALAT1* depletion in TPC1 cells is associated with increased mitochondrial respiration, potentially through indirect effects on nuclear-encoded mitochondrial pathways.

**Figure 6:**
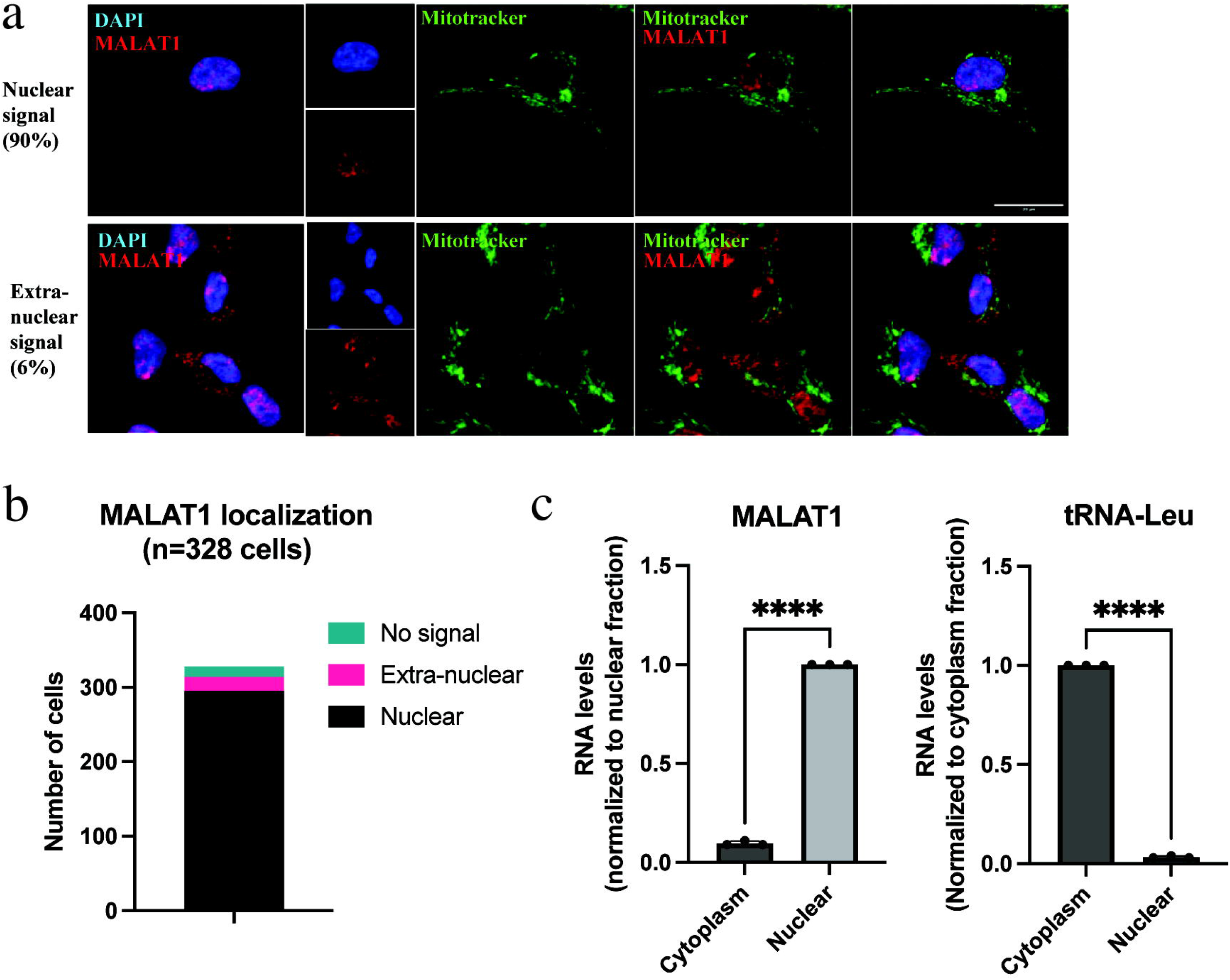
MALAT1 has a nuclear localization in TPC1 cells. (a, **b)** *MALAT1* (red) localization was analyzed by RNA-FISH combined with Mitotracker staining for mitochondria visualization. Representative images of exclusively nuclear *MALAT1* or cytosolic MALAT1 are shown (scale bar = 30 μm). Cell quantification from four biological replicate is shown in (b). **(c)** Nuclear and cytoplasmic extracts were prepared from TPC1 cells to quantify *MALAT1* levels by RT-qPCR. The tRNA^Leu^ was used to control the purity of fractions (n = 3 biological replicates, Welch’s t-test, ****: p<0.001).

## Discussion

In this study, we reported that lncRNA *MALAT1* levels were significantly higher in a cohort of Chilean patients with thyroid cancer with positive lymphatic infiltration. An exploratory normalized transcriptomic comparison of one high-risk tumor and its matched adjacent tissue additionally identified descriptive differences in cancer-related genes and biological processes. Using the papillary thyroid cancer cell line TPC1, we showed that *MALAT1* depletion was sufficient to reduce the proliferation and migration of cells, while reducing mitochondrial membrane potential and increasing mitochondrial respiration. Notably, we did not find evidence of *MALAT1* mitochondrial localization in these cells, suggesting regulation of genes at the nuclear level.

### MALAT1 as a diagnostic and prognostic marker in Chilean thyroid cancer samples

The potential of *MALAT1* as a biomarker for cancer diagnosis and prognosis has been extensively discussed in recent years (42). While meta-analyses have shown that *MALAT1* has moderate diagnostic performance, a considerable percentage of patients test negative (33% according to (43)), a trend that was also observed in our data. In contrast, *MALAT1* correlates more strongly with prognosis, as elevated levels have been linked to lower survival rates and increased metastasis risk (44), a pattern found in our small cohort. As current meta-analyses have notable limitations (including reliance on data primarily from American or Chinese patients, a focus on studies published in English or Chinese, and potential publication bias due to underreporting of negative results (43, 44)) our study contributes to addressing these gaps by providing data from a Chilean patient cohort.

Briefly, Chile has a population of 18.5 million and an admixed genetic background of European and Native American ancestry (45). Although our cohort was small, our results are consistent with previous *MALAT1* studies in thyroid cancer, which have examined from 10 to 195 tumor samples from different subtypes (19–23). In Chile, where papillary thyroid cancer is the most common subtype (2), our cohort of 16 patients included 14 papillary thyroid cancer cases. While we did not observe a consistent trend in *MALAT1* expression distinguishing tumor samples from adjacent tissues (**Fig. 1**), we found a significant association between *MALAT1* levels and lymphatic infiltration using both a paired estimation approach and multiple linear regression, even within this small cohort. These findings further support *MALAT1*’s role as a prognostic marker in Chilean thyroid cancer patients. Increasing the sample size in future studies will be essential to strengthen this conclusion. This association was not confirmed when we modeled *MALAT1* expression directly in a population-scale linear regression using the TCGA-THCA cohort. Lymph node status did not predict *MALAT1* expression once age, gender, and molecular subtype were accounted for. Interestingly, molecular subtype was itself a strong independent predictor (**Supplementary Figure S1**). This discrepancy between our clinical cohort and the larger, unselected TCGA-THCA population underscores the need for ecological observational studies to clarify *MALAT1*’s prognostic value.

### MALAT1 is involved in multiple processes in papillary thyroid cancer

Our exploratory transcriptomic comparison of a high-risk (ATA 3) thyroid tumor identified substantial differences in normalized gene abundance relative to matched adjacent tissue. Following normalization and conservative expression filtering, 4,407 genes showed at least a twofold higher normalized abundance and 2,399 showed at least a twofold lower normalized abundance in the tumor. Because the analysis was performed on a single matched pair, these values describe within-patient differences and should not be interpreted as statistically significant differential-expression calls.

Importantly, *MALAT1* was enriched in the tumor tissue (Patient 16). The difference in fold-change magnitude observed between RT-qPCR (1.85-fold) and RNA-seq (∼5-fold) may reflect differences in reference normalization, sub-optimal PCR amplification efficiency, or library composition effects. Crucially, alternative RNA-seq normalization methods (DESeq2 and TMM) produced closely concordant estimates of approximately fivefold higher abundance, supporting the directional robustness of this finding. Similarly, CDH2, and VIM showed higher normalized tumor abundance, whereas PCDH10 showed lower abundance (**Fig 2d**). These selected genes have previously been linked to *MALAT1* regulation in other experimental systems (46–48). However, the present RNA-seq analysis establishes only descriptive associations between *MALAT1* and these genes within a single patient and does not demonstrate direct regulation. Furthermore, because intact tumor and adjacent tissues were compared, differences in cellular composition between the two samples may contribute to both gene-level and pathway-level differences. The transcriptomic findings should therefore be considered hypothesis-generating and require validation in additional biological samples.

Previous studies have assessed transcriptional and phenotypic changes in thyroid cancer using various cell lines, including the anaplastic thyroid cancer lines SW1736, KAT18, and 8505C, the follicular subtype FTC133 (20–22), and the papillary subtype TPC1 (19). Notably, Zhang et al. (19) observed that EMT induction in TPC1 cells leads to *MALAT1* upregulation. Our findings further support the idea that *MALAT1* is not merely a byproduct of cancer progression but an active contributor, as its depletion in TPC1 cells reduces proliferation, alters cell cycle progression, and impairs migration (**Fig. 3**).

The association between *MALAT1* and epithelial-to-mesenchymal transition (EMT) is a well-documented, pan-cancer hallmark (16). Rather than positing a novel thyroid-specific driver role for the *MALAT1*–EMT axis itself, the primary contributions of our study are providing the first clinical evidence of this axis in a Hispanic/Chilean thyroid cancer cohort, and uncovering a regulatory layer governing how *MALAT1* link cancer cell fitness with cellular energetics

Although the specific molecular mechanisms underlying *MALAT1*’s effects remain beyond the scope of this study, previous research offers valuable insights. For instance, in lymphoma cells, *MALAT1* recruits the epigenetic silencing complex PRC2 to repress expression of cell cycle inhibitors *P21* and *P27* (49). A similar mechanism was described for *PCDH10* repression in gastric cancer (48). Furthermore, recent epigenomic analyses indicate that *MALAT1*-PRC2 complexes target and regulates hundreds of genes in breast cancer cells (50). In parallel, *MALAT1* functions as a competing endogenous RNA (ceRNA), sequestering miRNAs and preventing them from acting on their target mRNAs (18). For example, in lymphoma, *MALAT1* sponges miR-195, which regulates PD-L1 (a key modulator of T cell function) and subsequently affects *VIM* expression (51). A more direct mechanism was reported in hepatocellular carcinoma, where *MALAT1* sponges miR-30a-5p, a direct regulator of *VIM* (52). Additionally, in anaplastic and follicular thyroid cancer cell lines, *MALAT1* was found to promote proliferation and invasion by regulating *IQGAP1* expression (21). Whether similar mechanisms operate in papillary thyroid cancer requires further investigation.

### MALAT1 knockdown is associated with increased mitochondrial respiration

Finally, we explored the potential relationship between *MALAT1* and respiratory metabolism. Like many cancer cells, TPC1 cells exhibit high glycolytic activity (**Fig. 4b**), a characteristic that supports tumorigenesis, proliferation, and migration under both hypoxic and aerobic conditions (24). Upon *MALAT1* depletion, we observed a reduction in mitochondrial membrane potential accompanied by increase in respiration (**Figures 4a,b**). Given that *MALAT1*-silenced cells also exhibited reduced proliferation, we hypothesize that this shift toward increased respiration may accompany the less aggressive phenotype observed in our experiments, though direct functional causality remains to be established. This effect could occur in parallel with, or as a consequence of, *MALAT1*-driven transcriptional changes.

*MALAT1* has been linked to glucose and lipid metabolism across various models. In cardiomyocytes and endothelial cells, its expression increases under high-glucose conditions (53, 54). In hepatocellular carcinoma, *MALAT1* enhances glycolysis while suppressing gluconeogenesis through translational regulation of the metabolic transcription factor TCF7L2 (38). Additionally, *MALAT1* knock-down in hepatocellular carcinoma cells reduces glucose uptake and lipogenesis by downregulating lipid metabolism-related genes ( 55), a metabolic shift also observed in prostate cancer cells following *MALAT1* depletion (56).

In our TPC1 model, *MALAT1* depletion was associated with increased mitochondrial respiration without detectable changes in glycolysis. Unlike previous reports describing *MALAT1* localization in mitochondria (25, 26), we found no evidence of this in TPC1 cells. Instead our findings are consistent with an indirect, non-mitochondrial axis through which *MALAT1* may influence nuclear pathways associated with mitochondrial function. Consistent with this, population-scale correlation analyses in the TCGA-THCA cohort showed that *MALAT1* expression is largely decoupled from oxidative phosphorylation and glycolysis gene expression, but is more strongly (and negatively) associated with mitochondrial dynamics and mitophagy transcripts in node-positive tumors, pointing toward a role in mitochondrial quality control rather than direct bioenergetic regulation (**Fig. 5**). To the best of our knowledge, no link has been found between *MALAT1* and nuclear genes involved in mitochondrial dynamics and mitophagy and the precise mechanisms will be subjected of further investigation.

### Limitations of the study

Finally, several methodological constraints of our functional experiments should be considered. First, functional loss-of-function assays were conducted using a single antisense oligonucleotide (ASO) sequence within a single papillary thyroid cancer cell line model (TPC1). While TPC1 serves as a well-established model of papillary thyroid carcinoma, evaluating these phenotypes across additional cell line models (such as BCPAP or K1) will be essential to establish generalizability across diverse thyroid cancer genetic backgrounds. Furthermore, although our ASO produced robust and selective knockdown of *MALAT1*, future studies incorporating a second, non-overlapping ASO, alongside re-expression rescue experiments, will be important to completely exclude sequence-specific off-target effects and confirm absolute phenotypic dependency on *MALAT1*. Consequently, our *in vitro* observations provide a foundational framework linking *MALAT1* to mitochondrial energetics in thyroid cancer, which should be expanded upon in multi-model validation studies

## Conclusion

Our study provides evidence linking *MALAT1* levels with lymphatic infiltration in Chilean patients and supports a functional contribution of MALAT1 to proliferation, migration, and mitochondrial bioenergetics in papillary thyroid cancer cells. Importantly, *MALAT1* appears to influence metabolism primarily through nuclear-related mechanisms rather than direct mitochondrial localization. These findings highlight *MALAT1* as both a potential prognostic biomarker and a regulator of cancer cell biology, emphasizing the need for further studies in larger patient cohorts to validate its clinical utility and to elucidate the precise molecular pathways through which it acts.

## Acknowledgements

The work was funded by UNAB DI-08-24/APP (to M.T), ANID FONDECYT 1240853 (to R.A.) and 1251799 (to A.E.), Pew Advancement Award (to R.A.), UNAB DI-03-22/NUC (to A.E, P.C and R.A.), UNAB DI-03-24/IA (to A.E.), ANID FONDEQUIP EQM220115 (to A.E.). We also thank the Flow Cytometry Facility at the Institute of Biomedical Sciences, UNAB, for its critical contribution to this work. A preprint version of this work was deposited in bioRxiv (57).

## Data availability

The datasets generated and/or analysed during the current study are available in the GEO repository, accession number GSE315375. (Link: https://www.ncbi.nlm.nih.gov/geo/query/acc.cgi?acc=GSE315375). Other data are available on request from the authors.

## Author contributions

PC, AAE, and RA designed, supervised and funded the study. MTL, AM, MM, DL, GR performed experiments. GN performed the bioinformatic analyses. GR, LM, and AAE performed statistical analysis. RA wrote the manuscript. PC, AAE edited the manuscript.

## Use of Artificial Intelligence

Generative AI (Claude, Anthropic) was used to assist in producing the correlation-matrix figure (Fig. 5) and in improving the English grammar, clarity and fluency of the manuscript text during revision. All AI-assisted content was reviewed, verified, and approved by the corresponding authors, who take full responsibility for the accuracy and integrity of the manuscript.

## Supplementary table legends

**Supplementary Table S1.** Quantification of *MALAT1* and clinical parameters measured in this study (related to Figure 1 and Table 1)

**Supplementary Table S2.** Raw and normalized gene-level abundance values from the exploratory RNA-seq comparison of one matched tumor–adjacent thyroid tissue pair. Raw counts, DESeq2-normalized counts, descriptive log2 fold change calculated from DESeq2-normalized counts, TMM-normalized CPM values, TMM log2 fold changes, expression-filter status, and descriptive abundance classifications are provided.

**Supplementary Table S3.** Gene Ontology Biological Process over-representation analysis for genes with higher and lower normalized abundance in the tumor sample.

## Supplementary Figure legends

**Supplementary Figure S1.** Multiple linear regression of *MALAT1* expression (log2 RSEM-normalized) on lymph node status (N0 vs. N1), age, gender, and TCGA molecular subtype (BRAF-like as reference, compared with RAS-like and Other/Not Applicable) in the TCGA-THCA cohort, restricted to patients with complete covariate data (N = 419). Overall model fit: R = 0.254, R² = 0.065.

**Supplementary Figure S2** Full-length blots analyzed in this article (related to figure 4c)

