## Supplementary Figure S1 for "*MALAT1* Levels Are Elevated in Thyroid Tumors from Chilean Patients with Lymphatic Infiltration and Are Linked to Metabolic Reprogramming"

Multiple linear regression of *MALAT1* expression (log2 RSEM-normalized) on lymph node status (N0 vs. N1), age, gender, and TCGA molecular subtype (BRAF-like as reference, compared with RAS-like and Other/Not Applicable) in the TCGA-THCA cohort, restricted to patients with complete covariate data (N = 419). Overall model fit: R = 0.254, $R^2$ = 0.065.

| Predictor | Estimate | SE | t | p |
|---|---|---|---|---|
| **Intercept (BRAF-like, N0, female, reference)** | 12.221 | 0.153 | 79.625 | <.001 |
| **Lymph node status: N1+ vs. N0** | 0.099 | 0.089 | 1.113 | .266 |
| **Age** | -0.003 | 0.003 | -1.081 | .280 |
| **Gender: Male vs. Female** | 0.062 | 0.093 | 0.665 | .506 |
| **Molecular subtype: Other/NA vs. BRAF-like** | 0.400 | 0.104 | 3.836 | <.001 |
| **Molecular subtype: RAS-like vs. BRAF-like** | 0.473 | 0.110 | 4.308 | <.001 |
