## Supplementary figures and images for "*MALAT1* Levels Are Elevated in Thyroid Tumors from Chilean Patients with Lymphatic Infiltration and Are Linked to Metabolic Reprogramming"

### Supplementary Figure S2

#### Supplementary Figure S2

Full-length blots analyzed in this article (related to figure 4c)

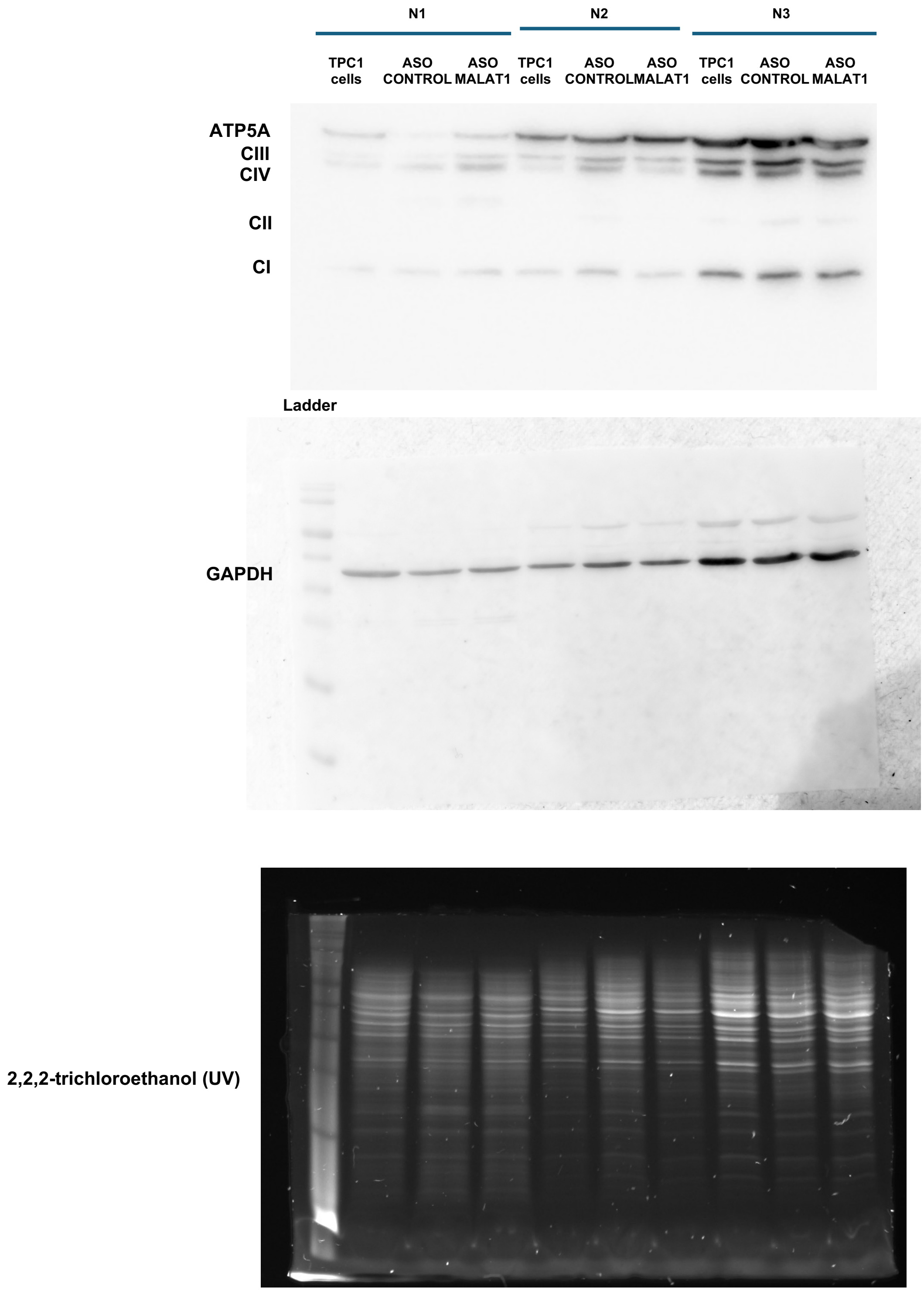
